# Linking dynamic photosynthetic traits to biomass: a functional data framework that pinpoints when photosynthesis limits growth

**DOI:** 10.64898/2026.11.03.763566

**Authors:** Junita Solin, Martin P. Boer, Djailey W. M. Gruijters, Fred A. van Eeuwijk, Tom P. J. M. Theeuwen

## Abstract

Photosynthesis responds rapidly to environmental changes, such as light fluctuations, and these responses affect how much carbon plants gain over time. Modelling genetic variation in photosynthesis over time opens a route for selection on photosynthetic performance. High-throughput chlorophyll fluorescence has enabled measurements of photosynthetic processes at high time resolution, yet methods to model the temporal and genetic variation in these time-series data remain scarce. Here, we combined splines and functional principal component analysis (FPCA) to model genotype-specific trajectories of fluorescence traits and capture their main variation in a few components. We applied this approach to 412 *Arabidopsis thaliana* MAGIC lines under daily sinusoidal and fluctuating light. The first component captured differences in overall trait level, whereas subsequent components captured differences in how trait values changed across days, mainly in the early growing period. These component scores predicted final dry weight reasonably well (average r = 0.72 under sinusoidal and 0.82 under fluctuating light). Information on changes across days added most predictive value when measurements were limited to a few days. Time-varying coefficients revealed that the associations between fluorescence traits and dry weight changed over time. For most traits, these associations were strongest early in the growing period. Our framework thus shows not only which photosynthetic processes are associated with biomass, but also when. This insight will point to the period in which photosynthesis limits growth and to the traits beneficial to select for.

## 1 Introduction

Photosynthesis is a central process for plant growth, but its rapid responses to environmental changes, such as fluctuating irradiance (Kaiser et al., 2018), make it difficult to phenotype and to select for. Due to this complexity, breeders and growers rank photosynthesis as a low priority for selection (CAPITALISE Consortium, 2024), and it has seen little direct improvement through breeding (Long et al., 2015). Historical yield gains have instead been attributed largely to selection on phenology and crop architecture (Welcker et al., 2022). Opportunities to breed for photosynthesis may therefore arise once we can phenotype it reliably and relate its dynamic behaviour to biomass accumulation or yield.

An increasing number of studies have shown the presence of natural genetic variation in photosynthesis (Acevedo-Siaca et al., 2020; Driever et al., 2014; Gao et al., 2024; Keller et al., 2022; Theeuwen et al., 2022). However, the strength of its relationship with yield varies considerably across studies. Some examples include moderate-to-strong correlations between photosynthesis and yield, for example, in wheat spike photosynthesis (*r* = 0.57; Molero & Reynolds, 2020), and in soybean (*r* = 0.80; Lopez et al., 2019). Other studies, however, reported weak associations (Driever et al., 2014, Keller et al., 2024). Notably, these studies typically relate yield to measurements that do not capture how photosynthesis changes over time, motivating approaches that do account for photosynthesis dynamics.

Advances in high-throughput chlorophyll fluorescence enable measurements of photosynthetic processes over time under dynamic environmental conditions (Baker, 2008; Cruz et al., 2016; Murchie & Lawson, 2013). Measurements at minute-scale resolution over the growing cycle capture rapid photosynthetic responses that influence biomass and yield. Studies that measured chlorophyll fluorescence under field conditions (Gao et al., 2024; Keller et al., 2024) showed that fluorescence-derived phenotypes are highly variable and changing with light intensity and developmental stages. Relating photosynthesis to biomass therefore requires repeated measurements both within the day and across the growing period. This, in turn, requires frameworks capable of integrating these dynamics over time and relating them to dry weight.

The most common framework to model dynamic photosynthesis is mechanistic modelling that integrates biochemical and plant physiological knowledge. Such a framework has been successful in scaling up leaf-level photosynthesis to quantify the impact of specific photosynthesis limitations on yield (Wu et al., 2019, 2023). However, mechanistic models were originally developed to describe processes, and not to model genetic variation. Recent work has shown that it is possible to link their parameters with genotypes by integrating crop growth models with genomic prediction (Jighly et al., 2023; Messina et al., 2023). For photosynthesis, however, such integration is less straightforward, as translating these measurements into genotype-specific model parameters requires assumptions about how fluorescence traits relate to the underlying processes. In addition, their reliance on numerous assumptions can introduce considerable uncertainty (Liu et al., 2025). On the other hand, highly flexible models from machine learning algorithms such as long short-term memory or transformers can indeed learn complex temporal patterns from the data, but they offer little mechanistic insight and are data-hungry (Hochreiter & Schmidhuber, 1997; Su et al., 2025; Van Houdt et al., 2020). Statistical modelling offers a middle ground, retaining interpretability while allowing greater flexibility than mechanistic models.

Functional data analysis (FDA) extends classical statistical methods by treating entire curves as the unit of analysis, rather than individual data points (Ramsay & Silverman, 2005). A common approach to reconstructing smooth curves from discrete time points is to use splines, which represent non-linear responses using piecewise polynomials. Adding a penalty on the roughness of the curve prevents overfitting, giving penalised splines (P-splines). P-splines have recently been implemented to model plant growth trajectories across genotypes and derive attributes that characterise genotypes (Pérez-Valencia et al., 2022; Pérez-Valencia et al., 2024). However, these attributes typically summarise global characteristics of the curves, while extracting localised features often requires prior knowledge of the expected patterns. Functional principal component analysis (FPCA) offers an alternative by identifying the main modes of variation in the data without requiring these patterns to be specified a priori (Ramsay & Silverman, 1997). These modes are represented by a few components that show how variation in a phenotype is distributed over time. Each genotypic curve is then assigned a functional principal component score for each component, indicating the direction and magnitude of its deviation from the mean curve. These scores can subsequently be used in downstream analyses to predict dry weight. Combining splines and FPCA, two tools from FDA, provides a framework for characterising genetic variation in photosynthetic time series and its association with dry weight over time.

In this study, we applied this framework to model genotype-specific trajectories of photosynthetic traits, use them to predict dry weight, and examine how their association with dry weight changes over time. Using a high-throughput phenotyping platform, we generated chlorophyll fluorescence time series data at 15-minute intervals for a MAGIC population of *Arabidopsis thaliana* under two contrasting light regimes: daily sinusoidal and fluctuating light. These relatively simple conditions provide a starting point, while the framework can be extended to more complex environments and to other crop species with broader genetic diversity. Specifically, we asked: (i) how genotypes differ in their photosynthetic trajectories within and across days; (ii) whether accounting for these trajectories improves the prediction of dry weight compared with daily means or individual time points; and (iii) which photosynthetic processes are associated with dry weight, and how these associations change over time.

## 2 Materials and methods

### 2.1 Plant material, experimental setup, and phenotyping

A total of 412 inbred lines of *A. thaliana* MAGIC panel (Kover et al., 2009) were grown in a controlled walk-in growth chamber at the Netherlands Plant Eco-phenotyping Centre (Fig. 1A). The phenotyping system PlantScreen^TM^ provided by Photon Systems Instruments (Drásov, Czech Republic) allowed us to simulate sunny and partly cloudy days, represented by daily sinusoidal and fluctuating light (Figure 1C). The total amount of light was similar for both regimes.

**Figure 1.**
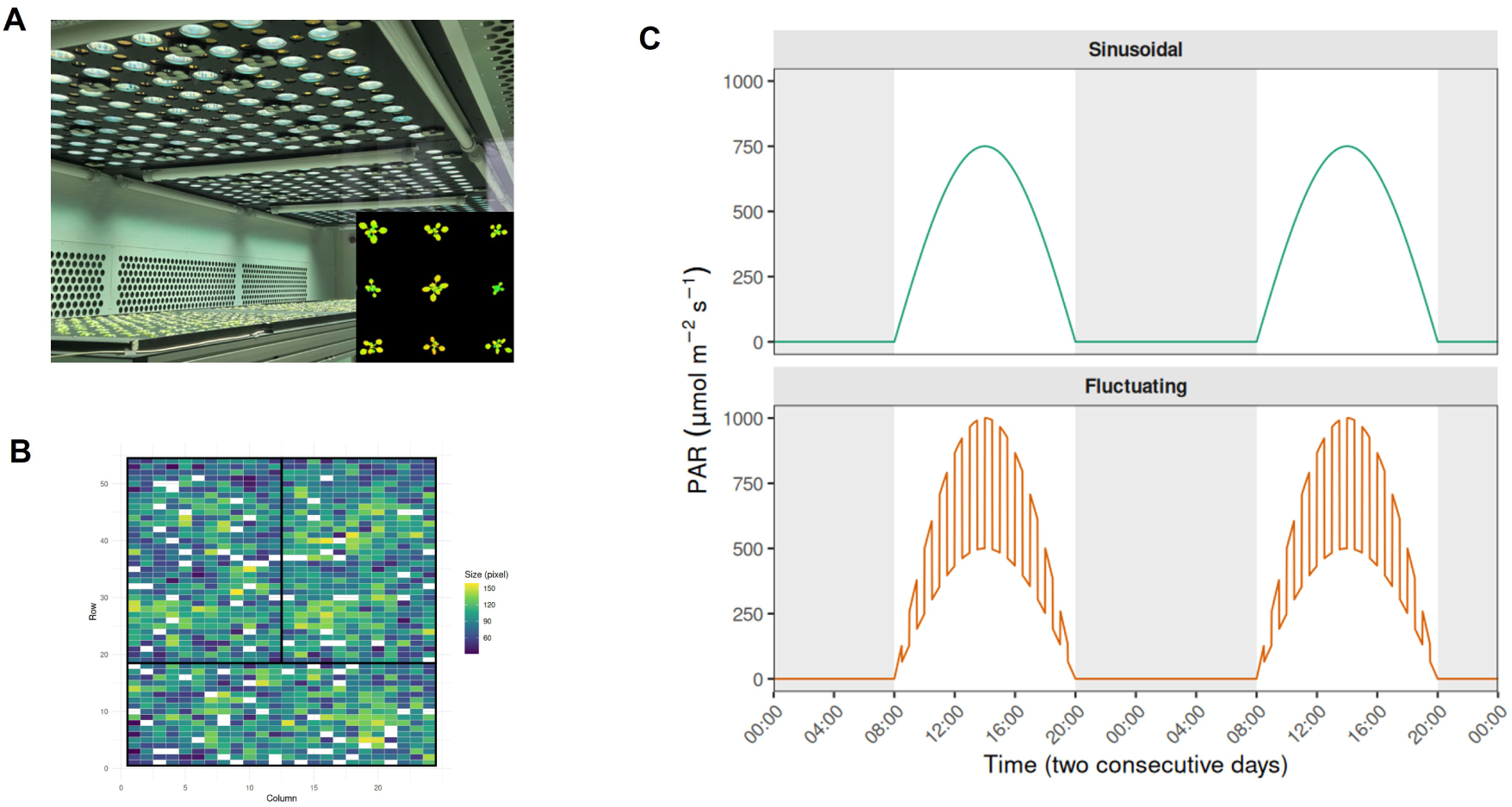
Experimental design, block layout, and lighting profiles. (a) A controlled-environment room equipped with fluorescence cameras enabling dense measurements at 15-min intervals under dynamically controlled growth conditions. An example of a fluorescence image is provided. (b) Experimental layout showing 412 genotypes across three complete blocks with 54 rows and 24 columns. (c) Photosynthetically active radiation (PAR) over two consecutive days under the daily sinusoidal and fluctuating regimes, with shaded regions indicating nighttime.

Seeds were germinated in Petri dishes on soaked filter paper with 0.65 mL demineralised water. Seeds were then placed in a dark, cold room (4 °C) to induce vernalization. After 5-6 days, these seeds were moved to a climate-controlled chamber (24 °C with 16/8 h day/night) for 24 h to induce germination. The germinated seeds were transplanted to the phenotyping facilities and sown on pre-soaked rockwool of Hyponex amended with MES buffer. The growth conditions were as follows: 12 h/12 h light/dark photoperiod, 480 ppm CO_2_, and 12°C/20°C temperature. The experimental layout was a randomised complete block design of three blocks, with a total of 1296 plants arranged in 54 rows and 24 columns.

Using chlorophyll fluorescence cameras installed at the ceiling, fluorescence traces were measured every 15 minutes, simultaneously on all the plants, starting from 6 days after sowing (DAS) up to 22 DAS, before plants started overlapping. A custom two-minute Pulse Amplitude Modulated (PAM) protocol was applied on dark- and light-adapted leaves to measure steady-state, maximum, and minimum fluorescence (Schreiber, 2004; Theeuwen et al., 2024). These values were used to derive 13 fluorescence traits that are listed in Table S1, following the equations in Baker, 2008; Kramer et al., 2004; Theeuwen et al., 2024. Outliers were defined based on the projected leaf area being lower than 1.5 times the interquartile range of the first quartile to remove plants that did not estalish well. Plants were removed from further analysis when outliers of projected leaf area were detected on more than three days. Negative values between 0 and -1 were set to zero, and values below -1 were set to NA. No additional filtering was performed to retain biological variation. The methods in this paper are summarised in Figure 2.

**Figure 2.**
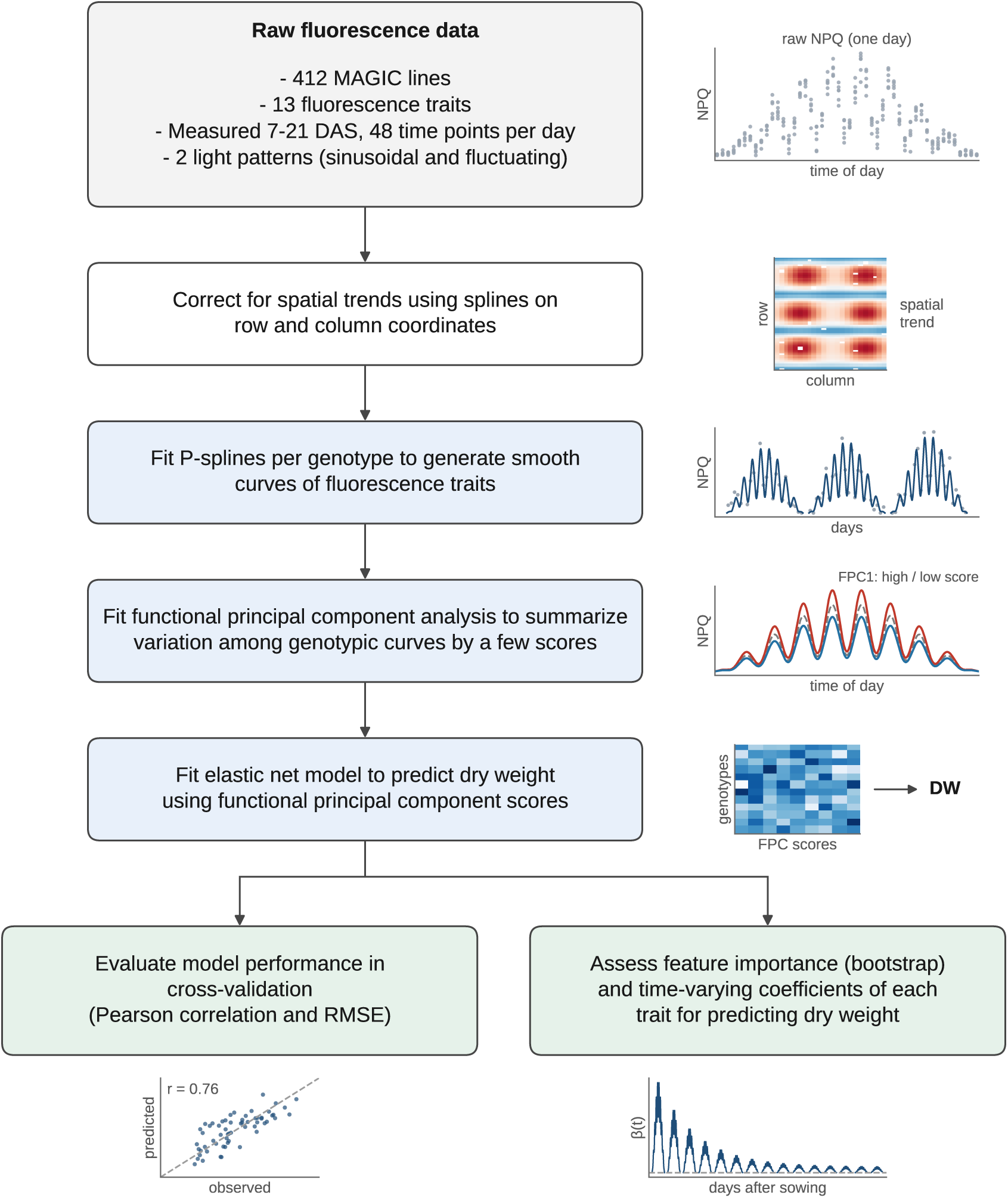
General flowchart of modelling time-series chlorophyll fluorescence data. Spatial correction was first applied to the fluorescence traits, followed by smoothing phenotypic curves using P-splines. Using singular value decomposition (SVD), we retained three principal components that capture the largest modes of variation among genotypic curves. Genotypic scores on those principal components were used as predictors in the elastic net model to predict dry weight under cross-validation. The model was evaluated using Pearson correlation and root mean squared error (RMSE). Feature importance was further analysed using the elastic net coefficients after bootstrapping. Considering the eigenfunctions, we could also assess the association of each fluorescence trait with dry weight over time (*β*(t)).

### 2.2 Spatial and design correction using SpATS

Spatial correction was applied to account for spatial trends in fluorescence traits, for example, due to the distance to the camera. Smoothing methods formulated within a mixed model were implemented to model spatial trends via the R package SpATS (Rodríguez-Álvarez et al., 2018) embedded in the R package statgenHTP. Row and column coordinates were used to compute two-dimensional smooth surfaces and treated as random factors on which the correction is based. The spatially corrected values were retained at the experimental unit level (plants). Spatial correction was applied separately to the 13 fluorescence-derived traits for each measurement time.

### 2.3 Modelling temporal genetic variability using splines

Splines are non-linear modelling techniques that use piecewise polynomials joined at specific points called knots. The domain of time is partitioned into intervals, and continuity constraints are imposed where the segments meet, which allows flexible and adaptive curves without assuming a fixed functional form (Hastie et al., 2009; Wood, 2017). Splines are well suited to model photosynthesis time-series data that follow complex diurnal and developmental dynamics. Such local dynamics would be ignored by simple parametric curves (e.g., a sinusoid). Splines can smooth measurement noise and convert discrete observations into a continuous curve which forms the basis of the subsequent functional analyses.

To begin with, a single-day model fitted separately for each genotype, with *p* indexing the 3–6 replicate plants within that genotype, is written as follows:

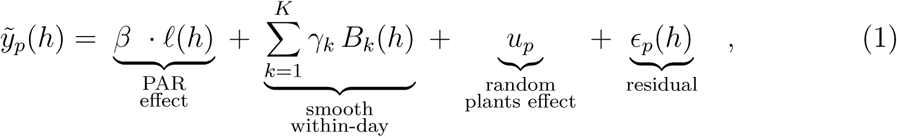

where *ℓ*(*h*) denotes PAR at within-day time *h* and 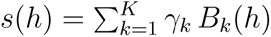 is a P-spline over within-day time, where *{B_k_*(*h*)*}* has a cubic B-spline basis and coefficients *γ_k_*. Smoothness is enforced by a second-order penalty on differences between adjacent coefficients, giving a penalised-spline (P-spline) formulation (Eilers & Marx, 1996) of which a relatively large number of knots can be used without overfitting.

In contrast to standard smoothing, our model decomposes each time series into different time scales: a smooth term over the time of day *h*, capturing the diurnal response, and a second smooth term over days after sowing *d*, capturing the developmental trend across days. In addition, the model includes the effect of light intensity and a random effect for individual plants, so that genotype curves are estimated while accounting for both the light regime and plant-to-plant variation:

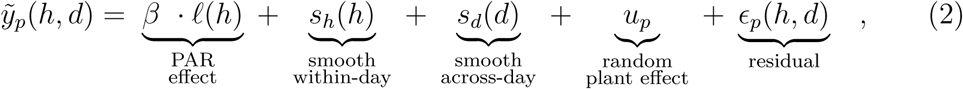

where *s_h_* and *s_d_* are P-splines with 40 and 10 knots, respectively, chosen relative to the number of time points within a day and the number of measurement days. The model was fitted separately for each genotype and trait using the R package LMMsolver (Boer, 2023), which represents the P-splines as a mixed model, so that the smoothing penalties are estimated by restricted maximum likelihood (REML) rather than chosen manually or via cross-validation.

### 2.4 Dimension reduction and curves decomposition via FPCA

After representing the photosynthesis time-series as smooth curves, we aimed to derive attributes that capture differences among genotypes. Global summaries, such as area under the curve, will overlook local dynamics. Conversely, spline coefficients are difficult to interpret, as each describes only a local part of the curve and their values depend on knot placement. Meaningful differences between genotypes, like differences in response timing, are distributed across many coefficients. To this end, functional principal component analysis (FPCA) was applied, as this method decomposes the variation among curves into a set of functional principal components (FPCs) (Ramsay & Silverman, 1997, 2005). These FPCs capture the main modes of variation among genotypic curves. Each genotype is then characterised by a few uncorrelated scores by projecting the deviation of its curve from the population mean onto the FPCs.

For each fluorescence trait, FPCA was applied to the set of *n_g_* genotype-specific smooth curves. Each curve *Z_i_*(*t*) (genotype *i*) on the full time domain *t* was decomposed as the mean curve plus a weighted sum of FPCs/eigenfunctions:

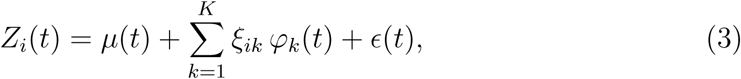

where *µ*(*t*) is the mean population curves, *φ_k_*(*t*) the *k*-th eigenfunction, *ξ_ik_* the score of genotype *i* on that component, and *K* the number of retained components. The scores were obtained by projecting the deviation of genotype *i* curve from the population onto the *k*-th FPCs. Eigenfunctions represent shared patterns that best explain how genotypic curves differ from each other.

We applied FPCA separately for each trait. Each smooth genotypic curve was evaluated on a fine, equally spaced grid spanning the observed time range, giving a matrix of genotypes *×* time points. We first computed the population mean curve *µ*(*t*) as the average of the genotypic curves at each time point, and subtracted it from each curve. A singular value decomposition was then applied to this centred matrix, **X***_c_* = **UDW**^T^, using the function fpca2s of the R package refund (Goldsmith et al., 2026; Xiao et al., 2016). The columns of **W** are the eigenfunctions *φ_k_* evaluated on the grid, and the entries of **UD** are the genotype-specific scores *ξ_ik_*. Because the curves had already been smoothed by the P-spline model and were evaluated on a dense grid, the eigenfunctions were not smoothed further. We retained the first *K* = 3 components per fluorescence trait, which on average explained 99% of the variation.

### 2.5 Dry weight predictions and feature importance via elastic net

#### Dry weight predictions

To relate the photosynthetic trajectories to biomass, the genotypic scores obtained from FPCA were used as predictors of final dry weight. For each of the 12 fluorescence traits with time-series measurements, the scores of the first three FPCs were used as predictors. Average Fv/Fm over time was used additionally, giving 37 predictors in total. We fit this model using elastic net regression, which balances a lasso penalty that shrinks the coefficients of uninformative predictors to zero with a ridge penalty that groups correlated predictors (Zou & Hastie, 2005). We used the R package `glmnet`, with predictors standardised internally, the mixing parameter fixed at *α* = 0.5, and the penalty parameter *λ* selected by 10-fold cross-validation (*λ*_min_). When *α* = 0.5, it gives an equal mix of ridge and lasso penalties, and *λ*_min_ is the value of the penalty chosen based on the lowest mean cross-validated error.

Reference models were proposed to evaluate the contribution of temporal variability of photosynthesis in predicting dry weight (Table 1). To robustly assess model performance, we applied 5-fold cross-validation repeated 100 times using Pearson correlation and root mean squared error (RMSE).

**Table 1.**
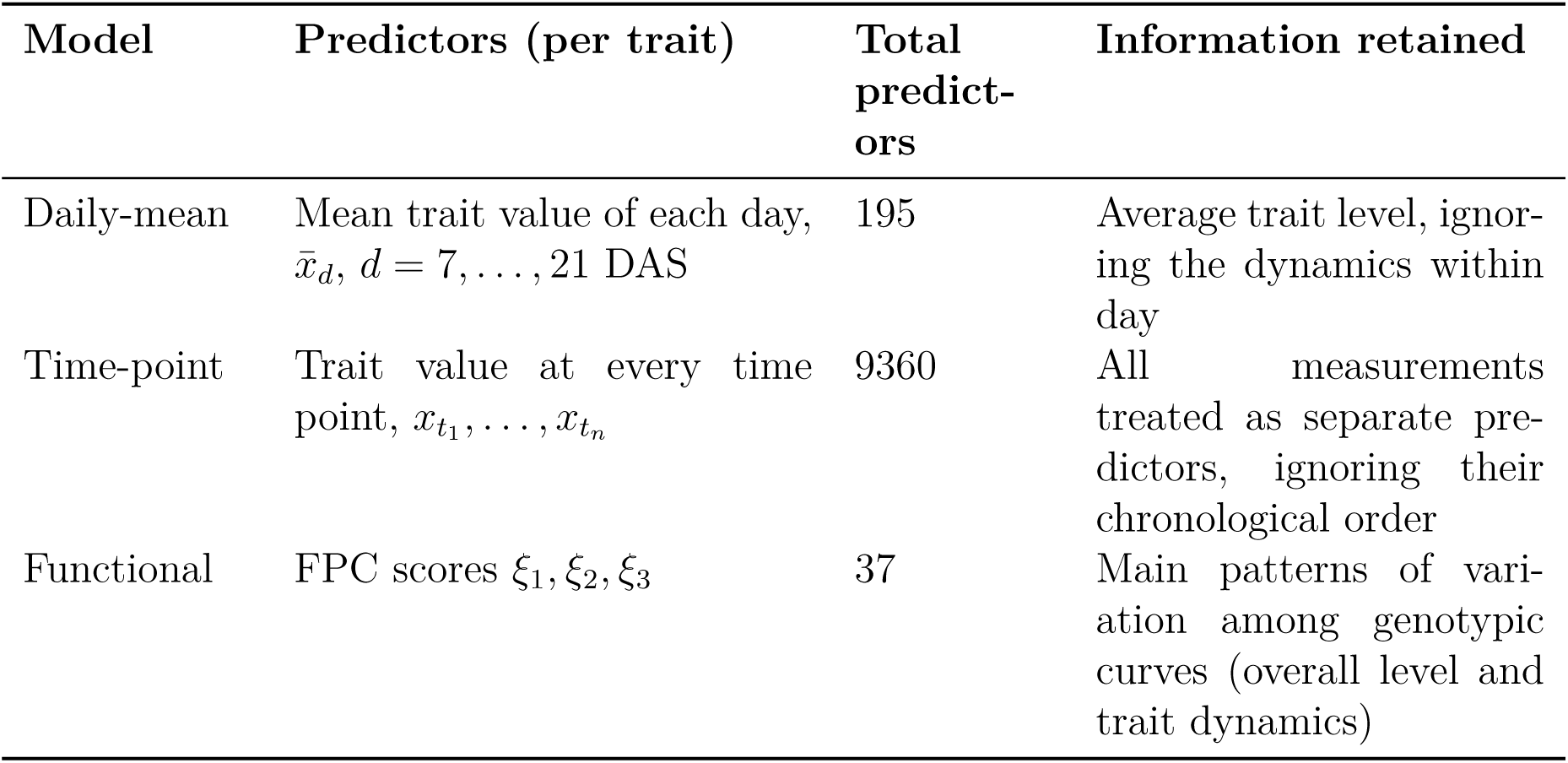
Models compared for predicting final dry weight (DW). Comparing the functional model with the daily-mean model tests whether temporal patterns add information beyond the daily trait level. Comparing it with the time-point model tests whether representing the time series as spline curves improves prediction accuracy rather than assuming every measurement as a separate predictor.

| **Model** | **Predictors (per trait)** | **Total predict-ors** | **Information retained** |
|---|---|---|---|
| Daily-mean | Mean trait value of each day, x̄<sub>d</sub>, d = 7, …, 21 DAS | 195 | Average trait level, ignor-ing the dynamics within day |
| Time-point | Trait value at every time point, x<sub>t<sub>1</sub></sub>, …, x<sub>t<sub>n</sub></sub> | 9360 | All measurements treated as separate pre-dictors, ignoring their chronological order |
| Functional | FPC scores ξ<sub>1</sub>, ξ<sub>2</sub>, ξ<sub>3</sub> | 37 | Main patterns of vari-ation among genotypic curves (overall level and trait dynamics) |

#### Feature importance

To assess which photosynthetic processes associate with dry weight, feature importance was assessed by fitting the elastic net model (*λ*_1se_) to 100 bootstrap resamples of the genotype panel. The parameter *λ*_1_*_se_* means the internal cross-validation error in the R package glmnet was set to be within one standard error of the minimum error. As a result, it produced a more parsimonious model than the absolute bestfitting model while performing nearly as well. This is also useful in assessing feature importance so that we retain the most robust contributing predictors.

In each resample, the fitted model predicts dry weight from the FPC scores of all traits,

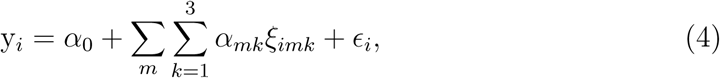

where *ξ_imk_* is the score of genotype *i* on the *k*-th functional principal component of trait *m*, and *α_mk_* the corresponding coefficient. The sum runs over the fluorescence traits measured as time series and Fv/Fm, which was measured once per day, entered the model as a single predictor and is omitted here for clarity. To compare features measured on different scales, coefficients were standardised as *α_mk_ ·* SD(*ξ_mk_*), where SD(*ξ_mk_*) is the standard deviation of the scores in the full data. For each feature, we computed its selection frequency, i.e. the proportion of resamples in which its coefficient was non-zero, and features selected in fewer than 50% of resamples were excluded.

The coefficients in Eq. (4) refer to FPC scores and therefore do not directly show when during the growing period a trait is associated with dry weight. They can be calculated back to the time domain using the projection of each genotypic FPC score onto the corresponding eigenfunction *φ_mk_*(*t*),

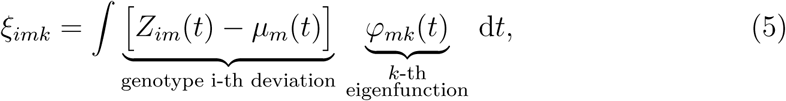

where *Z_im_*(*t*) is the spline curve of genotype *i* for trait *m* and *µ_m_*(*t*) the population mean curve. Substituting Eq. (5) into Eq. (4) gives

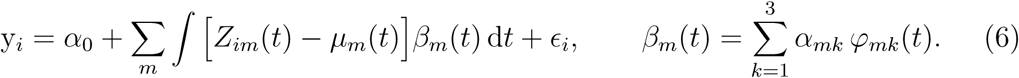

The coefficient *β_m_*(*t*) thus represents the contribution of trait *m* at each time point to the predicted dry weight. A positive value indicates that genotypes with above-average values at time *t* have a higher predicted dry weight, and a negative value indicates the opposite. We computed *β_m_*(*t*) from the coefficients averaged over the bootstrap resamples. Because all traits were included in the same multivariate model, *β_m_*(*t*) describes the association of trait *m* with dry weight conditional on the other traits.

## 3 Results

### 3.1 Splines captured within- and across-day dynamics of photosynthesis

The chlorophyll fluorescence imaging platform captured the rapid photosynthetic responses of *A. thaliana* genotypes as time-series measurements. To consider the temporal dependency of those data, we modelled each trait as a smooth curve using splines that allow each value to borrow information from neighbouring time points. Genotype-specific trajectories were a function of PAR and time at two scales: within-day and across-day using splines. Most fluorescence traits exhibited the light pattern of daily sinusoidal and fluctuating PAR. Such a pattern was less pronounced for basal dissipation (Φ_NO_) and the fraction of open photosystem II (qL) (Figure S1). Differences among genotypic curves of Col-0, Kn-0, and Rsch-4 were also observed in their within-day trajectories both under sinusoidal and fluctuating light (Figure 3A, 3B, and Figure S2). For NPQ, qE, and Φ_NPQ_, the largest differences tend to occur at midday under sinusoidal light and at the peaks of oscillation under fluctuating light. Meanwhile, Φ_II_ differed consistently throughout the day under sinusoidal light and under fluctuating light in the second half of the day. Other traits (i.e. Φ_NO_, qI, and qL) showed differences among genotypes that were irregular within a day. Fv/Fm is the theoretical maximum of photosystem II, which was measured once a day and therefore showed no within-day variation. These results show that splines capture the diurnal response of fluorescence traits and the differences in this response among genotypes.

**Figure 3.**
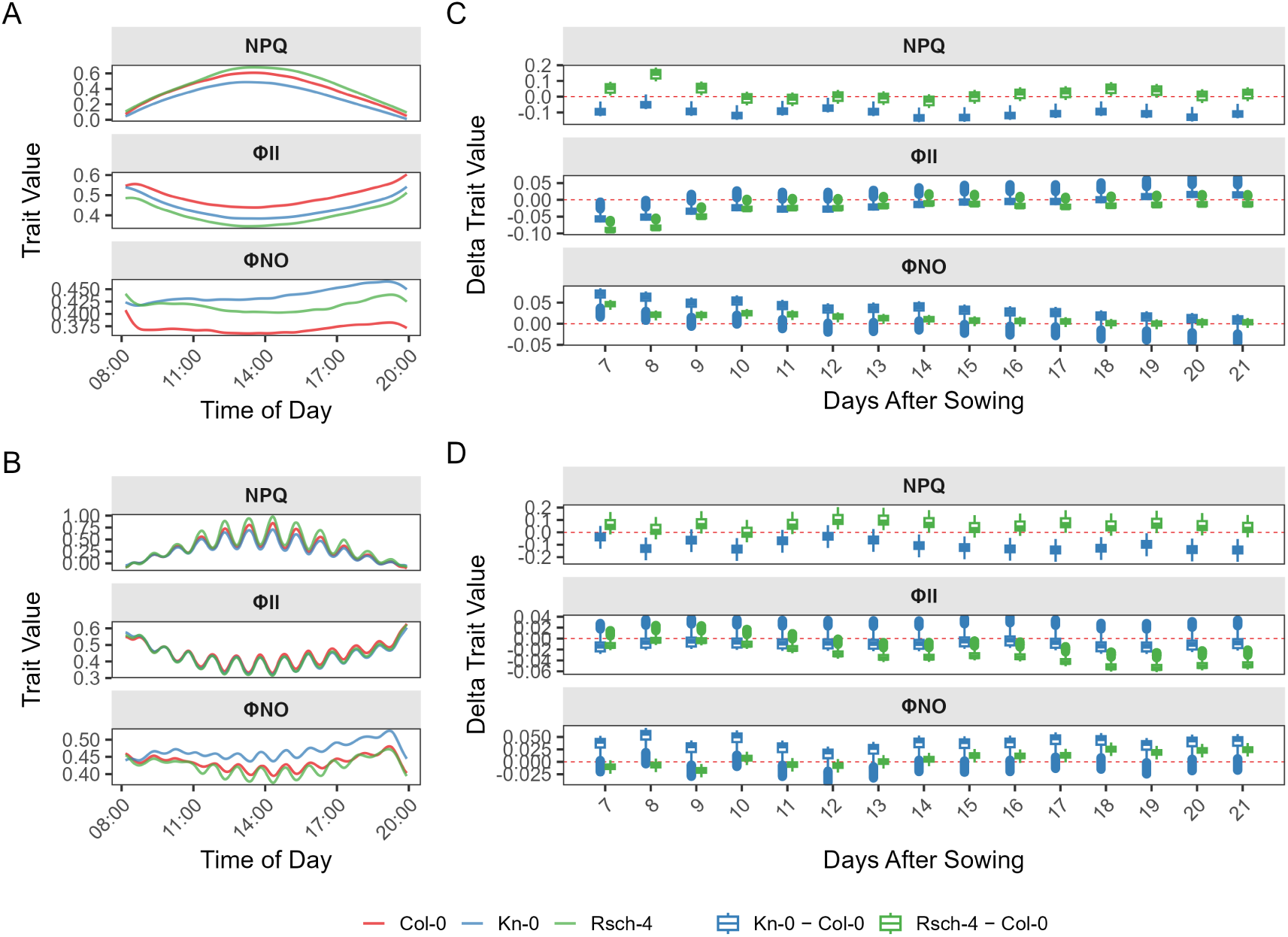
Genotype-specific curves vary within a day and across multiple days. (A) Spline curves of non-photochemical quenching (NPQ), basal dissipation (Φ_NO_), and efficiency of photosystem II (Φ_II_) shown at 7 days after sowing for three genotypes under sinusoidal light. For each genotype, the trait was modelled as a function of light intensity and time at two scales: within-day and across-day terms. (B) The fitted splines under fluctuating light. (C) Differences in trait values between Kn-0 and Col-0 (the reference) and between Rsch-4 and Col-0 plotted over days under sinusoidal light. These values were obtained after fitting splines on multiple days. (D) Differences in trait values between Kn-0 and Rsch-4 relative to Col-0 under fluctuating light.

To examine whether differences among genotypes were consistent from 7 to 21 DAS, we used three parental genotypes as an example and calculated the pairwise differences (delta) between Kn-0 and Col-0 and between Rsch-4 and Col-0. A positive delta indicates a higher trait value than Col-0, and a delta of zero indicates no difference in trait values with Col-0. For most pairs and traits, the delta remained consistently positive or negative throughout the growing period, indicating stable genotypic rankings over days (Figure 3C and 3D). However, the size of these differences changed over days. Under daily sinusoidal light, the differences of both Rsch-4 and Kn-0 from Col-0 in Φ_NO_ and Φ_II_ converged towards zero as the plants aged. For NPQ, this convergence was observed only for Rsch-4. Under fluctuating light, the difference in Φ_II_ increased over time, while that with Kn-0 remained stable. Adding a spline term for changes across days showed that genotypic differences in fluorescence trajectories occur not only within the day but also across days.

### 3.2 Temporal variability was dominated by genotypic mean differences

Genotype-specific spline curves revealed differences among genotypes both within and across days. To compare genotypes and relate their trajectories to dry weight, we aimed to summarise each curve into a small number of attributes. Using the spline coefficients directly is difficult, as each coefficient describes only a local part of the curve and depends on knot placement. Conversely, global summaries such as the area under the curve must be specified a priori and overlook local dynamics. We therefore applied functional principal component analysis (FPCA), which identifies a small number of orthogonal functional principal components (FPCs). These FPCs represent the main directions of variation among the genotypic curves. Projecting genotypic deviation from the population mean curve onto these FPCs yields a few genotype-specific scores, which we used to characterise genotypes.

The number of FPCs required depends on how much of the variability among curves they capture. For most traits, the first three FPCs explained, on average, 99% of the variability among genotypic curves. FPC1 accounted for most of this variability (on average 74% under daily sinusoidal light and 81% under fluctuating light), although its proportion varied among traits (Table 2). Traits related to non-photochemical quenching, such as NPQ, qI, and Φ_NPQ_, showed a lower percentage of variability captured by FPC1 than other traits. Traits describing similar processes, such as NPQ and NPQ(t) or qI and qI(t), differed in how variability was distributed across FPCs, which suggests differences in temporal variability. NPQ and qI were expressed relative to the dark-adapted Fm, which could vary among genotypes and over days, for example, when sustained quenching was still present at the time of measurement. In contrast, we calculated NPQ(t) and qI(t) only from light-adapted measurements, assuming a constant maximum PSII efficiency. Overall, FPC1 captured most of genotypic variability, whereas the contribution of FPC2 and FPC3 varied among traits and was largest for non-photochemical quenching traits, particularly qI.

**Table 2.** Percentage of variability among genotypic curves explained by each functional principal component.

|  | Sinusoidal light |  |  | Fluctuating light |  |  |
|---|---|---|---|---|---|---|
| Trait | FPC1 | FPC2 | FPC3 | FPC1 | FPC2 | FPC3 |
| Φ<sub>II</sub> | 88.6 | 8.0 | 3.4 | 87.1 | 10.0 | 2.8 |
| qL | 77.9 | 14.3 | 7.8 | 76.9 | 14.4 | 8.7 |
| NPQ | 69.3 | 18.5 | 12.2 | 75.0 | 15.6 | 9.3 |
| qE | 81.5 | 11.4 | 7.1 | 79.9 | 14.1 | 5.9 |
| Φ<sub>NPQ</sub> | 72.8 | 19.2 | 8.0 | 72.0 | 18.6 | 9.5 |
| NPQ(t) | 82.4 | 14.2 | 3.4 | 87.8 | 7.6 | 4.6 |
| qE(t) | 80.7 | 13.8 | 5.5 | 83.6 | 11.4 | 5.0 |
| Φ<sub>NPQ(t)</sub> | 89.6 | 7.2 | 3.1 | 89.5 | 8.0 | 2.5 |
| qI | 61.3 | 21.7 | 17.0 | 71.7 | 17.5 | 10.8 |
| qI(t) | 84.3 | 12.6 | 3.2 | 90.3 | 7.9 | 1.8 |
| Φ<sub>NO</sub> | 81.8 | 13.0 | 5.2 | 81.8 | 10.3 | 7.9 |
| Φ<sub>NO(t)</sub> | 79.3 | 12.7 | 8.0 | 74.5 | 20.7 | 4.8 |

To interpret each FPC, we reconstructed curves of hypothetical genotypes with scores two standard deviations above and below the population mean (hereafter high- and low-scoring genotypes). We use NPQ as the example here because it is a well-studied photoprotective trait with a strong and immediate response to light, so its curves show the within-day dynamics most clearly. Within the day, a genotype scoring high on FPC1 had higher-than-average NPQ around midday under sinusoidal light, and at the peaks of the light fluctuations under fluctuating light (Figure 4A, B). To examine these patterns over the growing period (7–21 days after sowing/DAS), we plotted the variance among genotypes explained by each FPC over days (Figure 4C, D). The variance explained by FPC1 was relatively stable across days under daily sinusoidal and fluctuating light, and its scores were strongly correlated with the average NPQ over days (average r = 0.99), confirming that FPC1 represents differences in the overall NPQ level among genotypes (or genotypic means) within a day. FPC2 and FPC3, which are independent of FPC1, captured how genotypes differed across days. FPC2 mainly distinguished genotypes in the early period (around 7-9 DAS). FPC3 captured a temporary deviation in the middle of the period under sinusoidal light (between 8 and 14 DAS), whereas under fluctuating light this pattern was less clear. These findings show that FPC1 captured overall trait level, whereas FPC2 and FPC3 captured differences in how NPQ changed over the growing period, mainly in the early and middle days, respectively. For other fluorescence traits, FPC1 similarly corresponded to the overall trait level, as shown by its strong correlation with the average trait value over days (average r = 0.99 under both light conditions). However, the shape represented by FPC1 varied among traits (Figure S4, S5). For Φ_II_, Φ_NO_, qI, and qL, high- and low-scoring genotypes on FPC1 differed in their trait level throughout the day, whereas for NPQ, qE, and Φ_NPQ_, the difference was concentrated around midday under sinusoidal light and at the light peaks under fluctuating light. Over the growing period, the variance captured by FPC1 remained the largest, although its value could fluctuate (increasing, such as Φ_II_ or decreasing, such as qE). In contrast, the variance of FPC2 and FPC3 decreased over days and reduced to zero as the plants aged. The exceptions were qI, qI(t), and qL, for which the variance of FPC2 followed a U-shaped trend, increasing again towards 21 DAS. In summary, differences in the curve shapes across days occurred mainly in the early days, while the later period genotypes mainly differed in their overall trait level.

**Figure 4.**
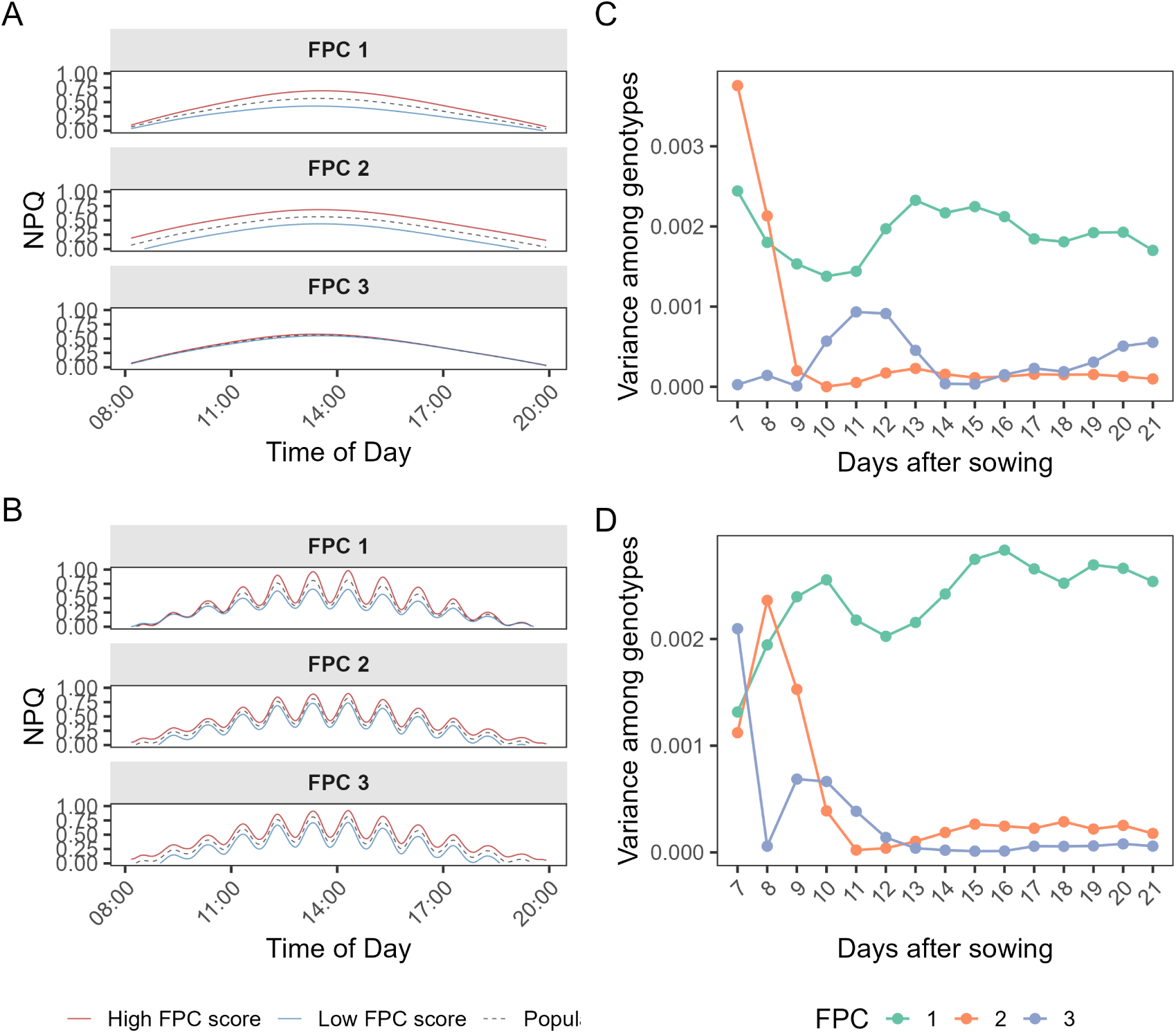
FPCA decomposes the within- and across-day dynamics captured by the spline curves into fewer components. (A) FPCA was applied separately per trait. The plots show NPQ at 7 days after sowing under sinusoidal light (A) and fluctuating light (B). Three components were retained, in total explaining 99% of the total variation, and each FPC corresponds to a distinct temporal pattern. The dashed curve shows the population mean, which is identical across all three FPC panels. The solid lines show the projected trajectories of genotypes with high or low scores on each component. (C) The variance of genotypic curves when projected on each FPC is shown per day from 7 to 21 days after sowing. This plot is for data under sinusoidal light, while the fluctuating light is shown at D.

### 3.3 Temporal dynamics predicted dry weight with fewer interpretable features

To relate the full trajectories of fluorescence traits to biomass, we obtained the genotypic scores from FPCA and used them to predict dry weight. We refer to this model as the functional model (37 predictors). For comparison, we used two reference models: a daily-mean model, using the average trait value of each day from 7 to 21 DAS as predictors (195 predictors), and a time-point model, using trait values at all time points as predictors (9360 predictors). All three models were fitted with the same elastic net procedure. Improved predictions by the functional model relative to the references indicated that the temporal dynamics of fluorescence traits were predictive of dry weight beyond their overall trait level. This model also allowed us to further investigate when during the growing period this relationship between photosynthetic traits and dry weight is strongest.

We could predict dry weight from fluorescence traits reasonably well, with overall predictions being more accurate under fluctuating light (average r = 0.72 and 0.82 for the functional model under daily sinusoidal and fluctuating light, respectively). Under fluctuating light, the functional model gave a higher correlation than the reference models (average r = 0.77 for the daily-mean and r = 0.80 for the time-point model), whereas under daily sinusoidal light it was on par with the reference models (Figure 5A). However, the RMSE did not show a consistent advantage for using the functional model. Under fluctuating light, its average RMSE was slightly higher than that of the time-point model (3.13 against 2.86 mg), and likewise under daily sinusoidal light, where it also marginally exceeded that of the daily-mean model (3.56 against 3.48 mg; Figure 5B). Relative to the average dry weight (16.31 mg under fluctuating and 20.07 mg under daily sinusoidal light), these differences between models correspond to less than 2% of the mean. These results suggest that much of the information used to predict dry weight is already captured by the average trait value of each genotype over time, which is shared by the functional model (via FPC1) and the reference models.

**Figure 5.**
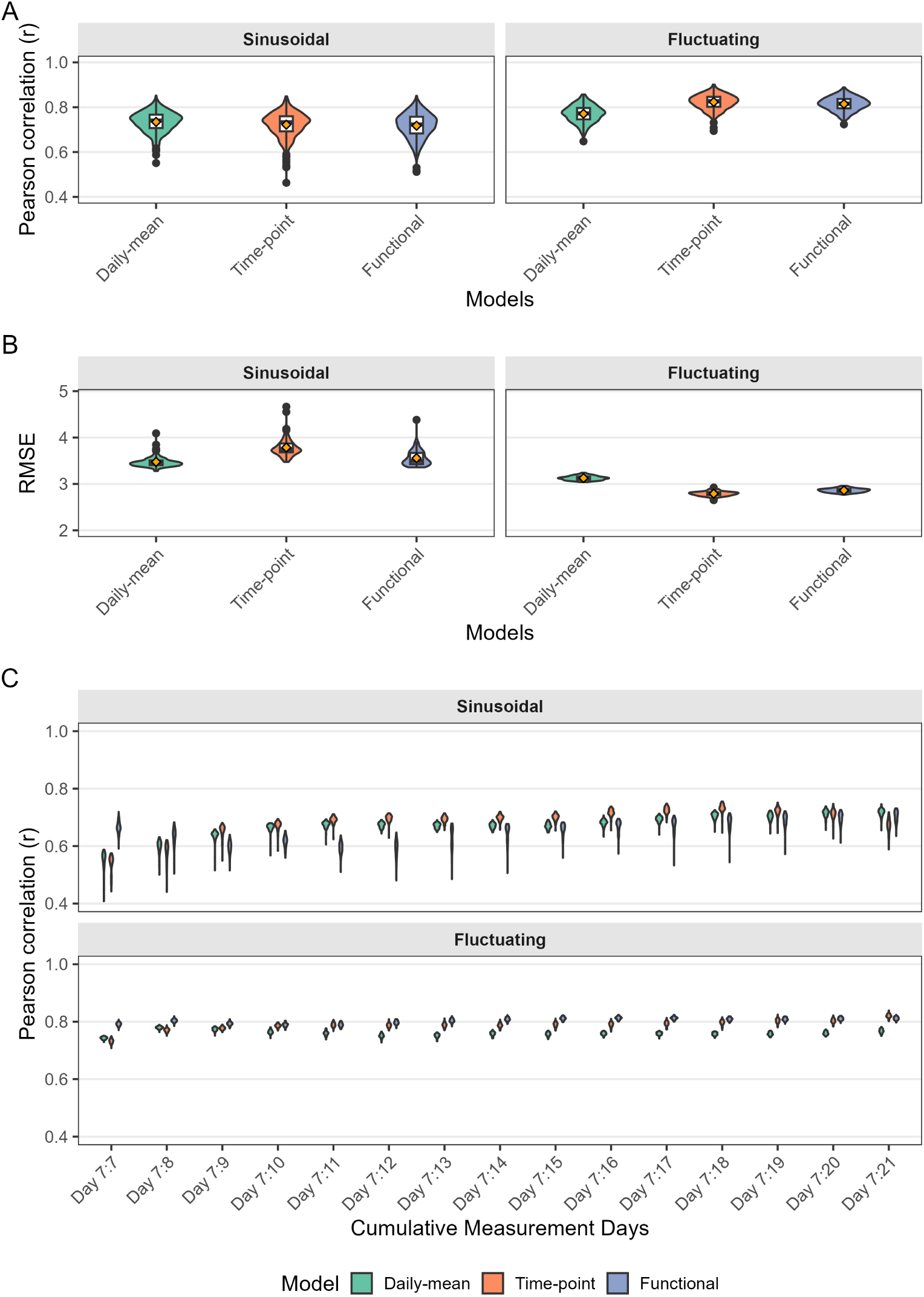
Adding temporal information improves dry weight prediction marginally, more evidently when measurements are restricted to early days. (A) The functional model was compared against two models that did not incorporate temporal patterns (daily-mean and time-point approaches). Pearson correlation is shown as violin plots, computed from 5-fold cross-validation repeated 100 times. (B) Root mean squared error (RMSE) was computed as a complementary loss metric. For reference, RMSE can be compared against the average dry weight of 20.07 mg (daily sinusoidal) and 16.31 mg (fluctuating light). (C) Pearson correlation was calculated from time-series data, with measurements increasing cumulatively by one day. Violin plots show the distribution across 100 repeats of 5-fold cross-validation.

To examine when temporal dynamics contribute most to dry weight prediction, we compared the models using an increasing number of measurement days, from a single day (7 DAS) up to the full period (7–21 DAS). The functional model clearly outperformed the reference models when only the first days were included (Figure 5C). At 7 DAS, the difference was largest under sinusoidal light (r = 0.67 for the functional model, against 0.55 and 0.54 for the daily-mean and time-point models). The difference also appeared under fluctuating light, though less substantial (r = 0.79, compared with 0.74 and 0.73). As more days were added and the daily light pattern was repeated, the differences among models gradually decreased. With measurements from 7 to 10 DAS, the functional model performed similarly to the daily-mean model under sinusoidal light and to the time-point model under fluctuating light. This is consistent with the FPCA results, in which FPC2 and FPC3 captured genotypic differences in the shape of the trajectories mainly in the early days. Including photosynthesis dynamics provided the greatest added value when the number of measurement days was limited (the early period) and when light patterns were less repetitive.

### 3.4 Specific photosynthetic processes linked to dry weight depend on the light environment

Beyond predicting dry weight, the functional model allows us to identify which fluorescence traits and their dynamics contribute most to dry weight predictions. To this end, we examined the coefficients of each FPC of each fluorescence trait in the functional model across bootstrap resamples. Overall, fewer fluorescence traits and their dynamics contributed to predictions under daily sinusoidal light (Figure 6). Those traits were specific to the dynamics across days of photoinhibitory quenching (FPC2 of qI and FPC3 of qI(t)) and the overall trait level of energy-dependent quenching (FPC1 of qE(t)). Basal dissipation traits were also contributing, though with less magnitude, and the least contributing process was photochemistry. Under daily fluctuating light, more processes were contributing, with the most pronounced ones being the overall level and dynamics of photoprotection traits (FPC2 of Φ_NPQ_ and FPC1 NPQ). Photochemistry traits and their dynamics (FPC1 and FPC2 of qL) were also predictive. In summary, dry weight prediction were driven by both the overall level of fluorescence traits (FPC1) and their dynamics (FPC2 and FPC3). Under sinusoidal light, these traits were mainly related to photoprotection and photoinhibition, but under fluctuating light, the contributing traits included those related to photochemistry and basal dissipation.

**Figure 6.**
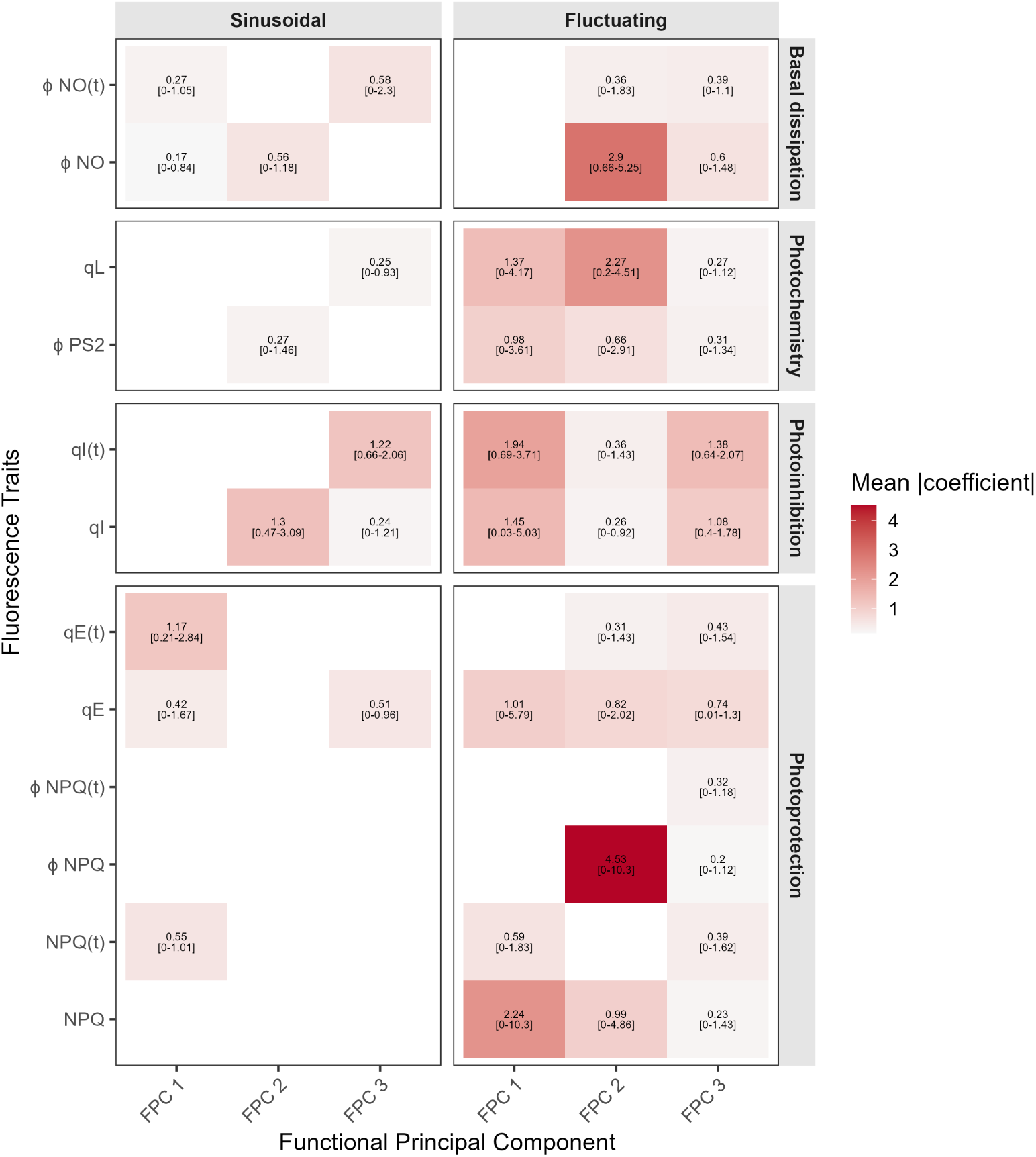
Fluorescence traits and their distinct temporal patterns contributed differently to predicting final dry weight. Values shown are mean absolute standardised coefficients across 100 bootstrap resamples with selected frequency > 0.5. Confidence intervals with 95% percentile are reported in brackets. Four categories group fluorescence traits by the photosynthetic process each is associated with. Larger values indicate a stronger contribution to prediction. Smaller values do not necessarily indicate an unimportant feature, as the elastic net model shares coefficients among collinear traits.

**Figure 7.**
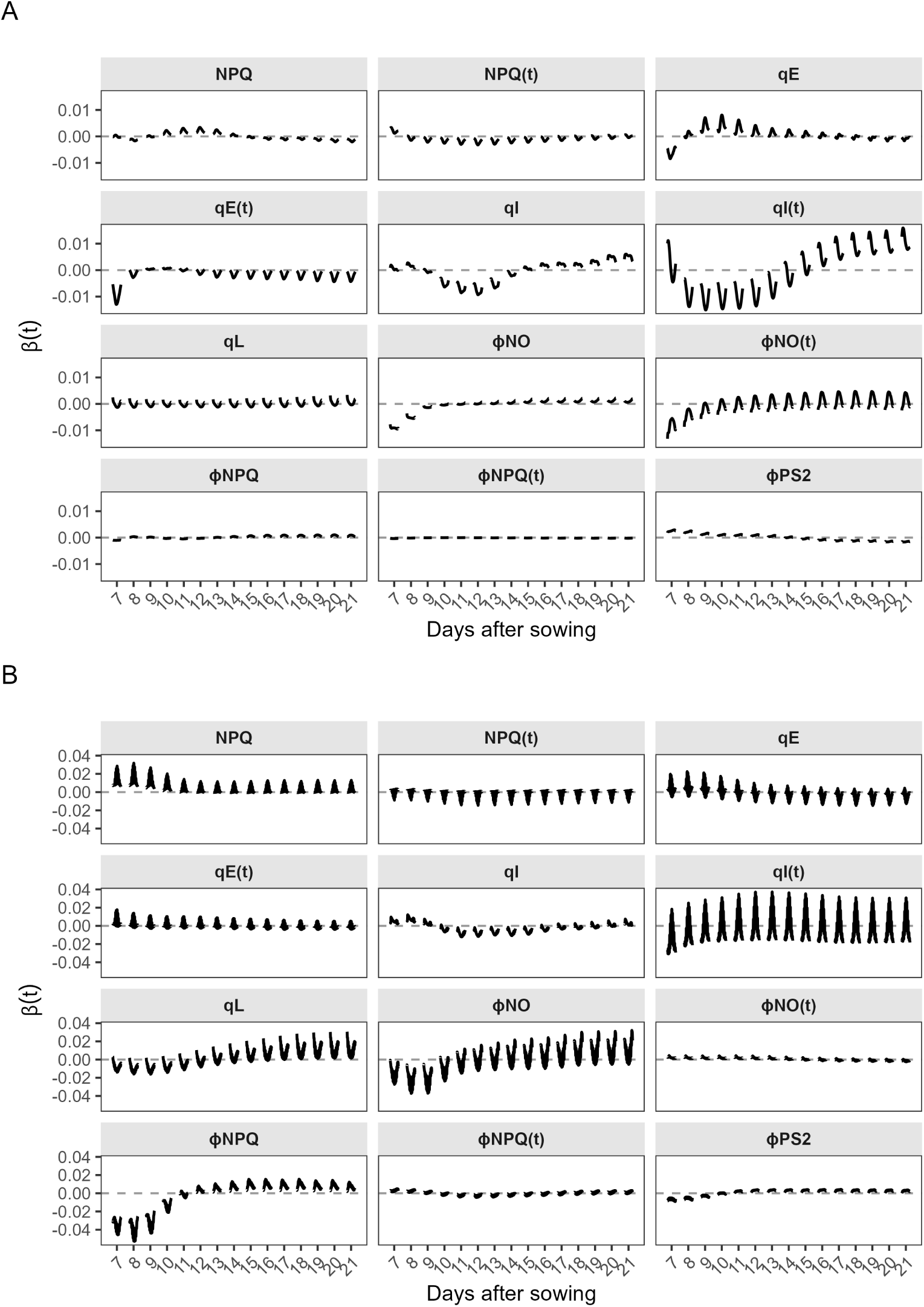
The relation between fluorescence traits and dry weight varies over time. A) Time-resolved coefficient functions *β*(*t*) under daily sinusoidal light. These values were computed as the sum of three FPCs of the eigenfunctions weighted by their elastic net coefficients. Zeros indicate no association with dry weight, and positive values indicate time points at which higher trait values are associated with greater dry weight. B) Time-resolved functions under fluctuating light.

To understand how the association of each fluorescence trait and dry weight changed over time, we calculated its time-varying coefficient *β*(*t*) (see Eq. (6)). Under daily sinusoidal light, the associations of photoinhibitory traits (qI and qI(t)) over days shifted from predominantly negative during the early-mid period to positive towards 21 DAS. Within a day, these associations were most negative around midday, coinciding with the peak of light intensity. In contrast, the associations of qE, Φ_NO_ and Φ_NO(t)_ with dry weight weakened over days. Under fluctuating light, the associations of the photoprotection-related traits NPQ and Φ_NPQ_ were most pronounced in the early period and close to zero later on, and within the day, they followed the light pattern with the strongest values around midday. Photochemistry-related traits (qL and Φ_II_), on the other hand, became gradually more positively associated with dry weight as the plants aged. These results exemplify that the time-varying coefficients enable us to resolve when, during the day and over the growing period, each photosynthetic trait is associated with biomass. For most traits they were strongest early in the growing period and weakened over time, while their within-day pattern depended on the light regime.

## 4 Discussion

In this study, we integrated time as an additional dimension in relating photosynthesis and biomass accumulation. We used splines to generate smooth trajectories of fluorescence traits for each genotype. Genotypic scores obtained from FPCA were used to summarise curve differences among genotypes. Finally, we predicted dry weight using those scores and obtained reasonable accuracy. Overall trait level (or genotypic means) contained a large part of the information that was associated with dry weight. We also successfully identified photosynthetic processes that drove the predictions over time. Previous studies have related photosynthesis to biomass under steady-state or fluctuating light, but without modelling the time aspect. Our approach explicitly modelled this relationship over time, which provides insight into which and when photosynthetic traits are associated with biomass.

Photosynthesis responds continuously to its environment, and accounting for the growth conditions under which it is measured is necessary. Previous studies on photosynthesis under fluctuating light proposed genotypic slopes describing how Φ_II_responds to light intensity (Keller et al., 2022, 2024) to account for light fluctuations. However, this approach still ignores how these genotypic slopes change over time. We argue that modelling response over time is equally important. We approached it by representing each fluorescence trait as a smooth function of time at two scales: diurnal response within-day and response across days, while also including light intensity as a covariate. This approach separates the response to light fluctuations from time and retains the full trajectory of each trait for each genotype.

FPCA characterised that variation among genotypic curves/trajectories of fluorescence traits was dominated by differences in overall trait level (genotypic means). Under both light regimes, FPC1 was strongly correlated with the average trait value over time, while FPC2 and FPC3 captured subtle differences across days. For most traits, these higher-order components accounted for only a small proportion of the variability. Notably, traits related to non-photochemical quenching, such as NPQ and qI, were captured by FPC2 and FPC3 to a larger share of the variability than remaining traits. This suggests that, for these traits, variability among genotypes is not explained only by how much energy they dissipated, but also by how this dissipation changed across days.

Dry weight was predicted reasonably well from genotypic FPC scores, though the proposed model performed similarly to the reference models under sinusoidal light. The functional model performed slightly better under fluctuating light, plausibly because genotypic variability was more pronounced under this light regime. As the large part of the information relevant for dry weight is contained in the overall trait level/FPC1 (strongly correlated with the average trait values), the differences in model performances were relatively small. The added value of temporal information captured by FPC2 and FPC3 became more evident in the early period, when measurements were limited and light patterns were less repetitive. In that case, the subtle differences in curve shapes provided information that complemented the overall trait level. These results suggest that the benefit of modelling photosynthetic trajectories is largest when measurements are limited or redundant information across days is low. Such conditions are common in field phenotyping, where light is irregular and measurement windows are short, on which the advantage of splines and FPCA could be more pronounced.

Beyond prediction, our approach allowed us to examine how the associations between fluorescence traits and dry weight change within the day and over the growing period. Under both light conditions, photoprotection and photoinhibition were important for predicting dry weight, with a strong association in the early part of the growing period. This pattern is consistent with source-sink dynamics, where enhancing photosynthesis has been shown to drive the greatest influence in growth during the early developmental phase (Kirschbaum, 2011; Torralbo et al., 2025). As plants age, internal limitations such as a reduction in the electron transport rate or a decrease in intercellular CO2 concentration or sink capacity become limiting instead. Previous work also highlighted the importance of rapid photoprotective responses for maintaining productivity under light fluctuations (De Souza et al., 2022; Kromdijk et al., 2016). Notably, photochemistry also emerged as an important predictor under fluctuating light, in particular the fraction of open Photosystem II (qL). Fluctuating light is a more challenging condition than daily sinusoidal light; therefore, higher efficiency in using light for photochemistry plausibly matters more. Previous studies show that genotypes with higher light use efficiency (the efficiency of converting the intercepted PPFR to photochemical energy) resulted in increased yield (Adachi et al., 2019; Keller et al., 2022, 2024). These condition- and time-specific associations could help define selection targets for improving photosynthesis from natural genetic variation.

Improvement in modelling the dynamics of photosynthesis and its relationship with biomass can be considered from two different areas: modelling and experiments. First, modelling using splines is beneficial in obtaining genotype-specific curves. However, fitting at the genotype level limits the partitioning of variability into genetic and non-genetic components. We could benefit from hierarchical modelling that includes genotype-to-plant structure in the splines (Di et al., 2009; Pérez-Valencia et al., 2022; Pérez-Valencia et al., 2024). We also observed the difficulties of splines in modelling edges of the time domain when there is no observation, such as at the beginning or the end of the day. Also, when handling irregularly sampled data that commonly occur in field measurements, the splines and FPCA implemented here would need to be extended.

Secondly, we realised that the experiment used simplified light conditions with a repetitive pattern. Although it provides tractable datasets for methodological development, light regimes with irregular fluctuations are closer to field conditions. Similarly, the study used *A. thaliana* plants due to their rosette-like morphology and fast growth. Testing our approach in other crops with more heterogeneous canopies will be a step closer toward assessing the relation between photosynthesis dynamics and yield relevant for breeding applications. Additionally, our measurements were based on chlorophyll fluorescence, which reflects photochemical efficiency but does not directly capture net carbon assimilation. When high-throughput techniques for measuring time-series carbon assimilation become available, applying our framework to such data would be a valuable extension. In summary, further studies are necessary to improve our modelling and experimental approach to linking photosynthesis dynamics and biomass, and this study serves as a starting point that shows time-series chlorophyll fluorescence is correlated with dry weight and identifies which photosynthetic processes drive it over time.

In conclusion, we propose a statistical framework that demonstrates the inclusion of the time dimension in understanding the relationship between photosynthesis and biomass across many genotypes. This approach provides a foundation for linking dynamic photosynthetic traits with crop yield under field conditions and pinpointing photosynthesis limitations. Moving forward, integrating this framework with genomics data will help identify the genetics underlying specific target traits for physiological breeding. Combining our statistical approach with mechanistic models would also facilitate scaling from the leaf to the canopy level when considering heterogeneous canopies (Ntakos et al., 2025; Wu et al., 2019, 2023). Ultimately, by extracting temporal insights from high-throughput time-series data, this framework bridges the long-standing gap between photosynthesis, a trait traditionally too complex to select for, and breeding efforts for enhanced yield and resilience.

## 5 Acknowledgements

We thank David Kramer for the discussion and feedback during the proposal writing. This publication is part of the project HARVEST of the research programme Genetics of Photosynthetic Limitations, which is (partly) financed by the Jan IngenHousz Institute.

## 6 Author contributions

J.S., T.P.J.M.T. and F.A.vE. conceptualised the study. D.W.M.G. conducted the experiments on the high-throughput phenotyping platform under the supervision of T.P.J.M.T., J.S., and T.P.J.M.T. curated the data. J.S. developed the statistical framework and performed the formal analysis, with M.P.B. contributing to the technical implementation of the splines. T.P.J.M.T, M.P.B., and F.A.vE. supervised the project. J.S. wrote the original draft. J.S., M.P.B., F.A.vE, and T.P.J.M.T contributed to reviewing and editing the manuscript. All authors approved the final version.

## 7 Data and code availability

The processed fluorescence data are available from the corresponding author upon reasonable request. All analysis code is available at <https://github.com/jsolin26/time-series-photosynthesis>..

## 8 Use of generative AI

The author used Claude to improve the language and readability of the manuscript and assist with code. The authors reviewed and edited the output and thus take full responsibility for the content of the publication.

## Supplementary Table

**Table S1.**
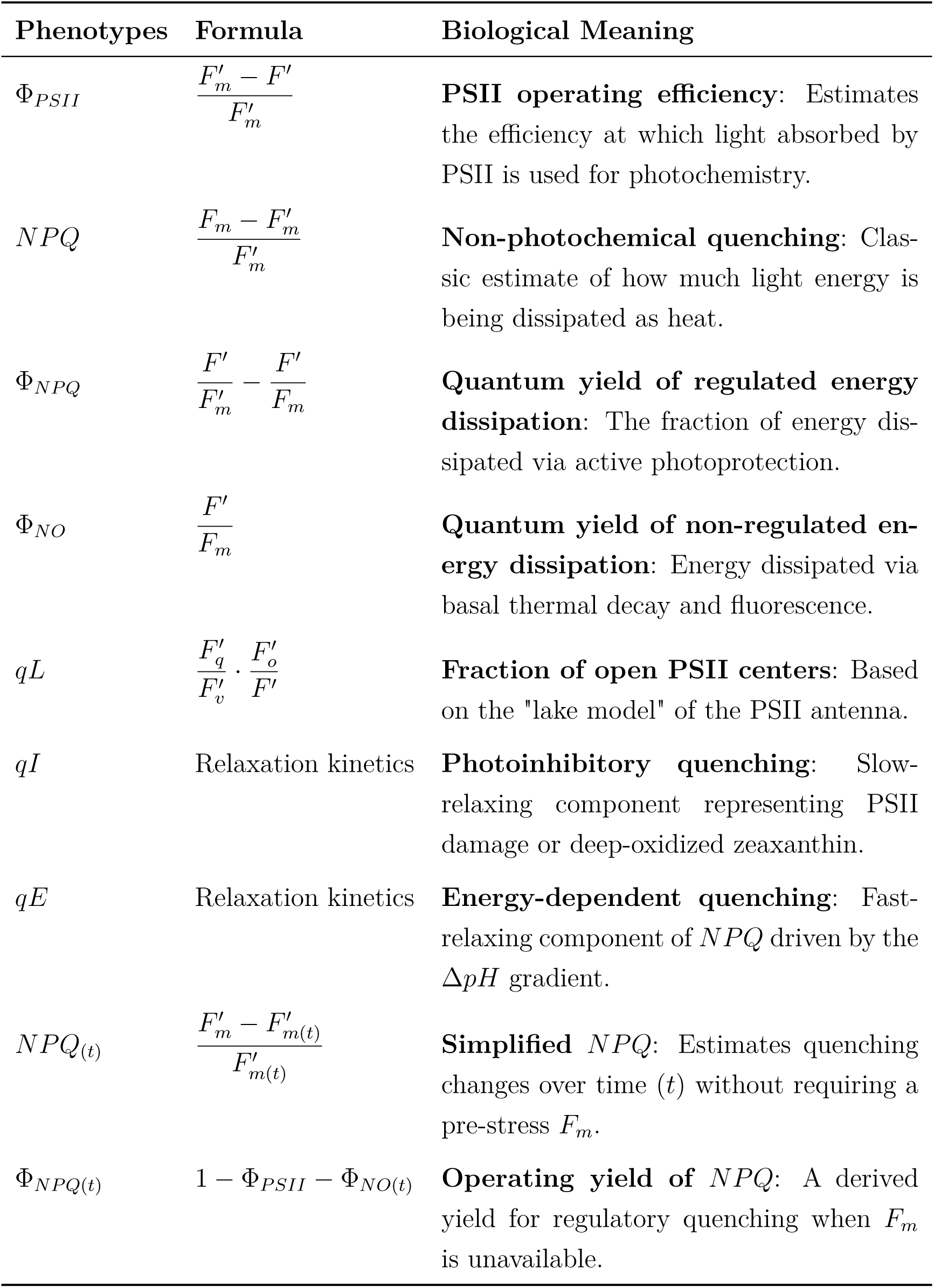
Extended Summary of Derived Photosynthetic Phenotypes from Chlorophyll Fluorescence Data and Biological Meanings.

| Phenotypes | Formula | Biological Meaning |
|---|---|---|
| Φ<sub>*PSII*</sub> | $\frac{F'_m - F'}{F'_m}$ | **PSII operating efficiency**: Estimates the efficiency at which light absorbed by PSII is used for photochemistry. |
| *NPQ* | $\frac{F_m - F'_m}{F'_m}$ | **Non-photochemical quenching**: Classic estimate of how much light energy is being dissipated as heat. |
| Φ<sub>*NPQ*</sub> | $\frac{F'}{F'_m} - \frac{F'}{F_m}$ | **Quantum yield of regulated energy dissipation**: The fraction of energy dissipated via active photoprotection. |
| Φ<sub>*NO*</sub> | $\frac{F'}{F_m}$ | **Quantum yield of non-regulated energy dissipation**: Energy dissipated via basal thermal decay and fluorescence. |
| *qL* | $\frac{F'_q}{F'_v} \cdot \frac{F'_o}{F'}$ | **Fraction of open PSII centers**: Based on the "lake model" of the PSII antenna. |
| *qI* | Relaxation kinetics | **Photoinhibitory quenching**: Slow-relaxing component representing PSII damage or deep-oxidized zeaxanthin. |
| *qE* | Relaxation kinetics | **Energy-dependent quenching**: Fast-relaxing component of *NPQ* driven by the Δ*pH* gradient. |
| *NPQ*<sub>(*t*)</sub> | $\frac{F'_m - F'_{m(t)}}{F'_{m(t)}}$ | **Simplified *NPQ***: Estimates quenching changes over time (*t*) without requiring a pre-stress *F*<sub>*m*</sub>. |
| Φ<sub>*NPQ*(*t*)</sub> | 1 − Φ<sub>*PSII*</sub> − Φ<sub>*NO*(*t*)</sub> | **Operating yield of *NPQ***: A derived yield for regulatory quenching when *F*<sub>*m*</sub> is unavailable. |

## Supplementary Figures

**Figure S1.**
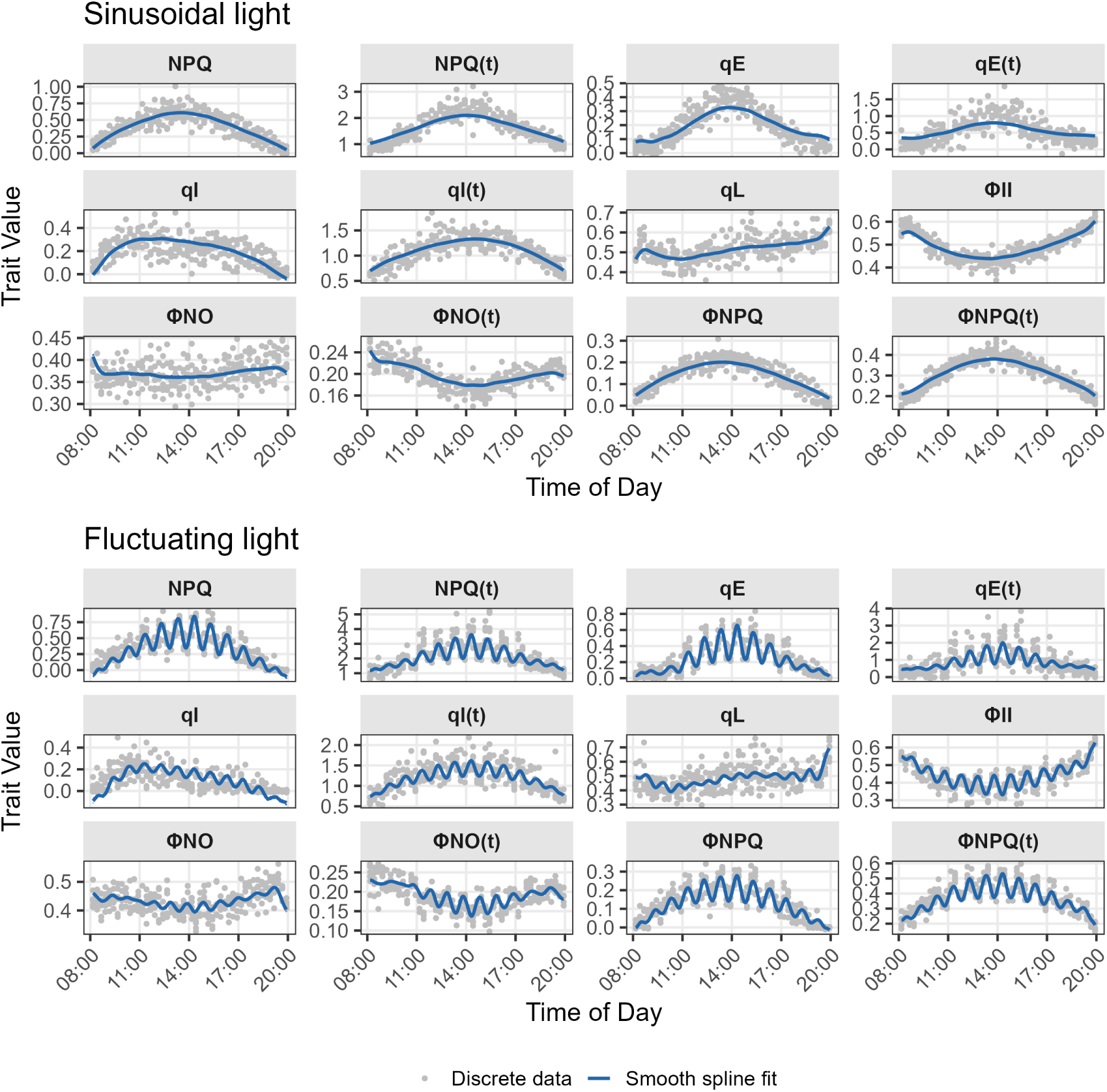
Spline fits on the discrete data of fluorescence traits at 7 days after sowing of Col-0. Smooth curves were generated by modelling light intensity and spline terms of time scale within-day and across days.

**Figure S2.**
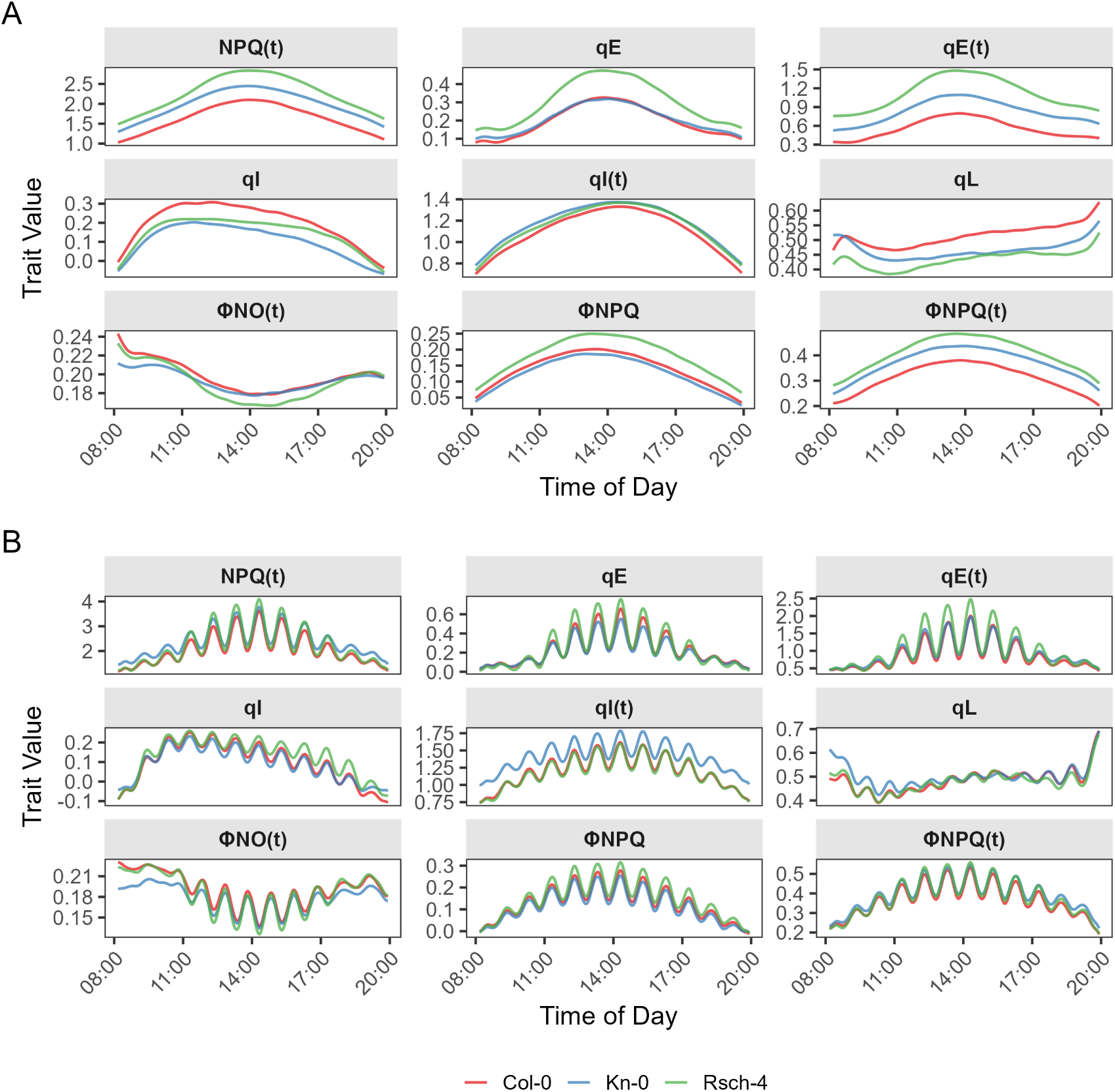
Genotype-specific spline fits for the remaining fluorescence traits shown at 7 days after sowing for three genotypes under light conditions: sinusoidal (A) and fluctuating light (B). For each genotype, the trait was modelled as a function of light intensity and time at two scales: within-day and across-day terms. We fitted splines independently.

**Figure S3.**
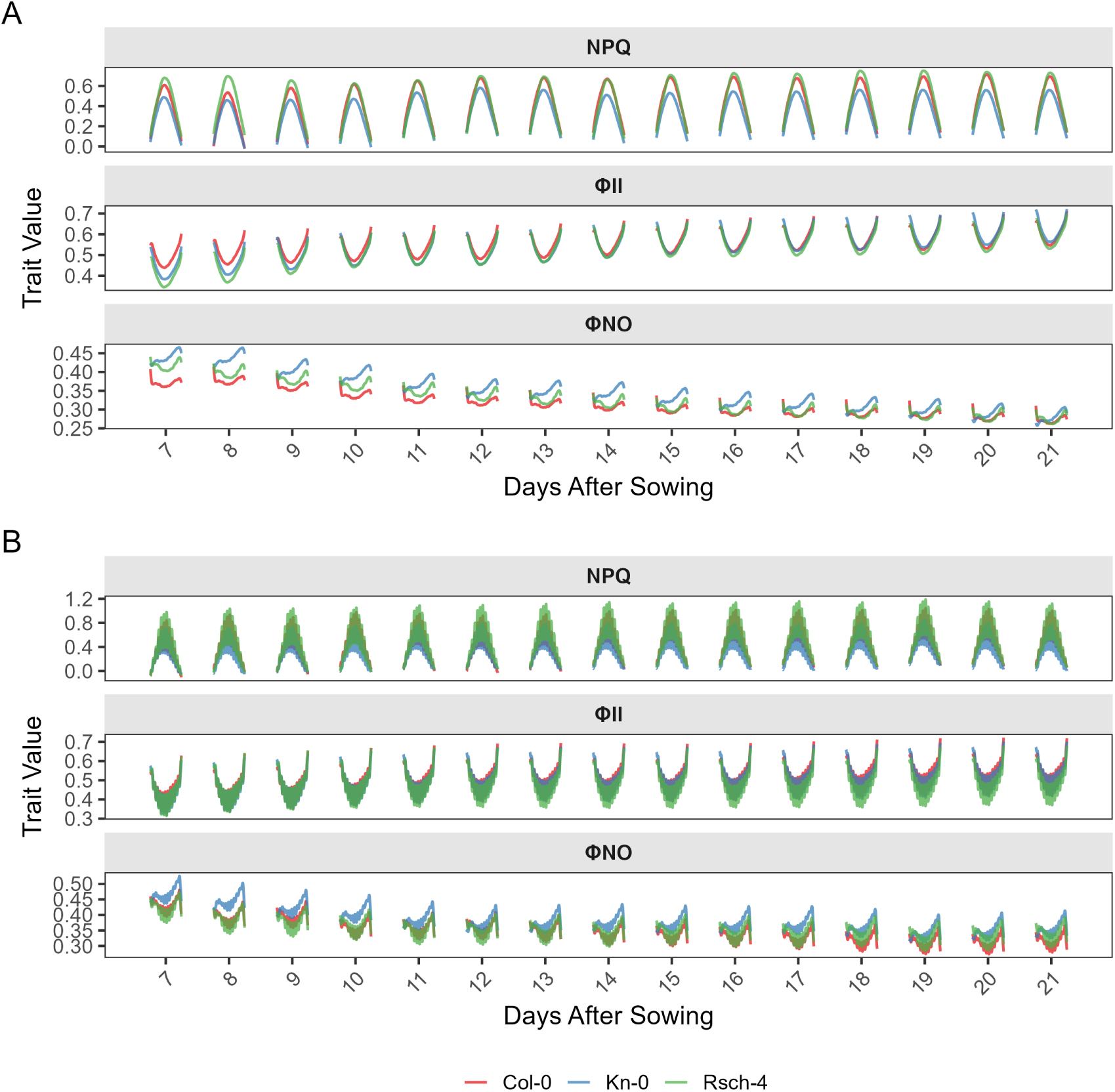
Spline fits of three genotypes shown for 7 to 21 days after sowing of non-photochemical quenching (NPQ), basal dissipation (Φ_NO_), and efficiency of photosystem II (Φ_II_). Two light conditions were applied: sinusoidal (A) and fluctuating light (B). For each genotype, the trait was modelled as a function of light intensity and time at two scales: within-day and across-day.

**Figure S4.**
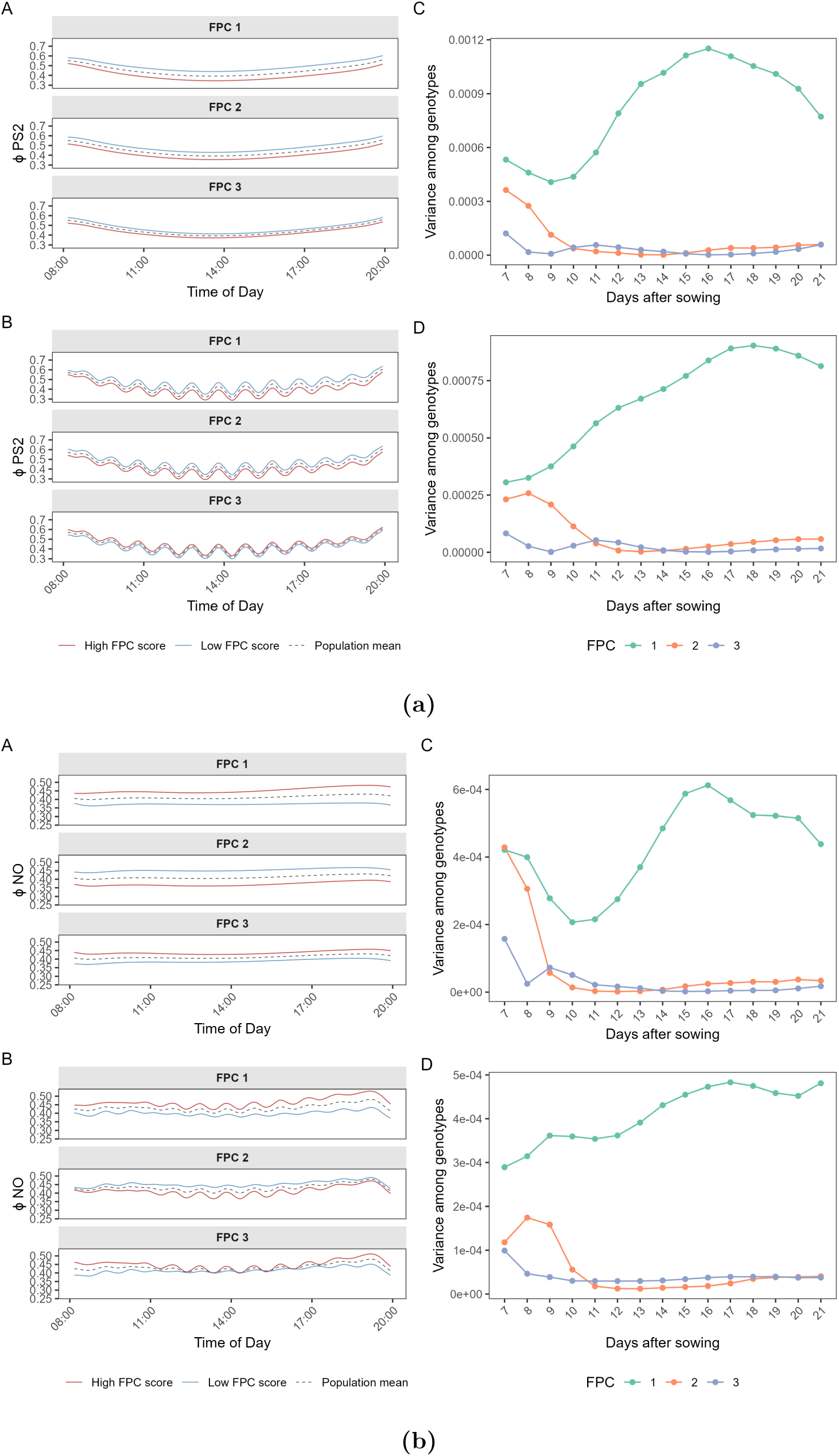
Functional principal components of (Φ_II_) (a) and (Φ_NO_) (b). For each FPC, three curves are plotted, where the dashed curve is the overall mean, identical across all three FPC panels for a given trait. The other two curves show the reconstructed

**Figure S5.**
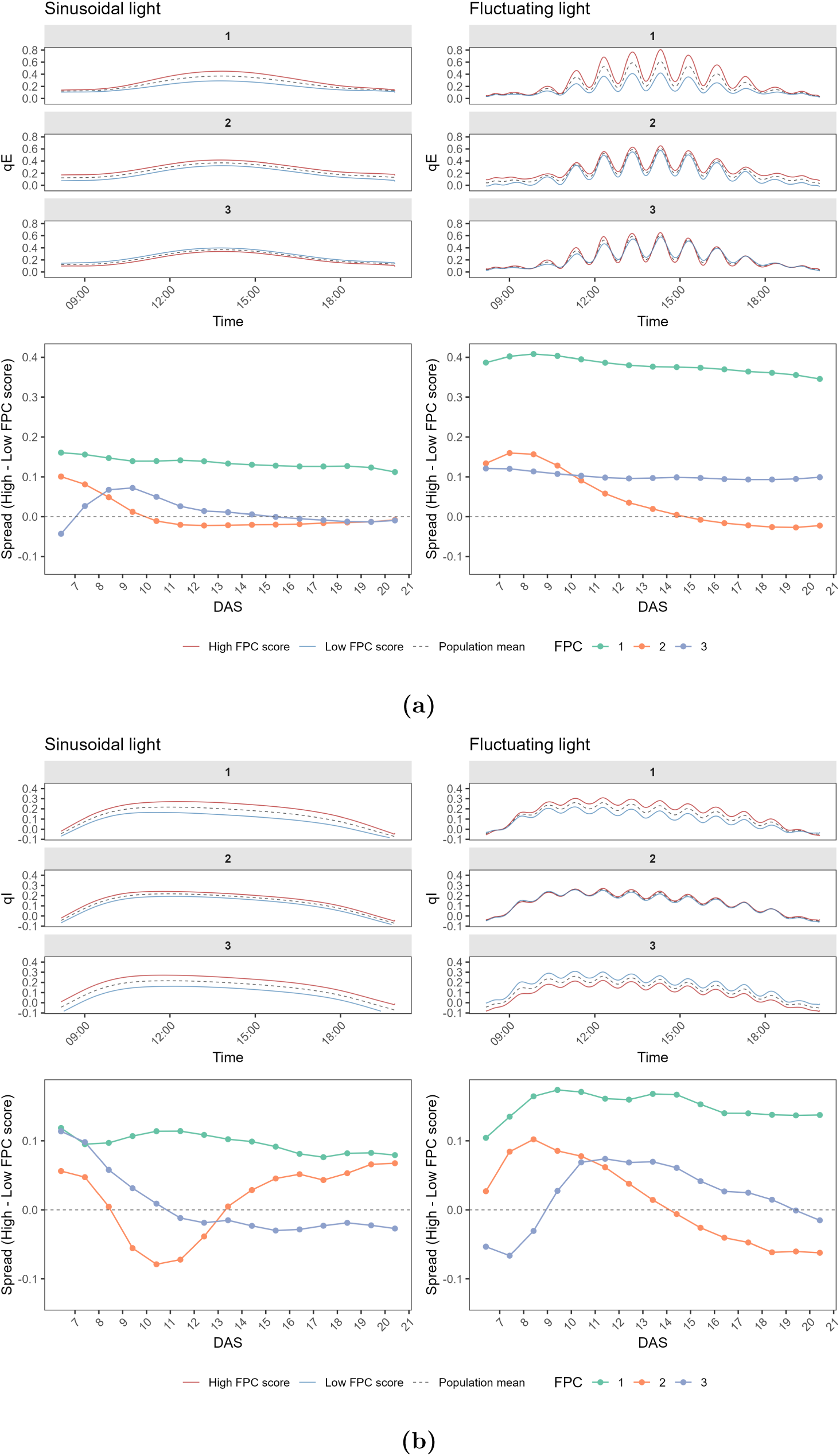
Functional principal components of qE (a) and qI (b) smooth curves. For each FPC, three curves are plotted, where the dashed curve is the overall mean, identical across all three FPC panels for a given trait. The other two curves show the projected

